# The ventral visual pathway represents animal appearance over animacy, unlike human behavior and deep neural networks

**DOI:** 10.1101/228932

**Authors:** Stefania Bracci, Ioannis Kalfas, Hans Op de Beeck

## Abstract

Recent studies showed agreement between how the human brain and neural networks represent objects, suggesting that we might start to understand the underlying computations. However, we know that the human brain is prone to biases at many perceptual and cognitive levels, often shaped by learning history and evolutionary constraints. Here we explore one such bias, namely the bias to perceive animacy, and used the performance of neural networks as a benchmark. We performed an fMRI study that dissociated object appearance (how an object looks like) from object category (animate or inanimate) by constructing a stimulus set that includes animate objects (e.g., a cow), typical inanimate objects (e.g., a mug), and, crucially, inanimate objects that look like the animate objects (e.g., a cow-mug). Behavioral judgments and deep neural networks categorized images mainly by animacy, setting all objects (lookalike and inanimate) apart from the animate ones. In contrast, activity patterns in ventral occipitotemporal cortex (VTC) were strongly biased towards object appearance: animals and lookalikes were similarly represented and separated from the inanimate objects. Furthermore, this bias interfered with proper object identification, such as failing to signal that a cow-mug is a mug. The bias in VTC to represent a lookalike as animate was even present when participants performed a task requiring them to report the lookalikes as inanimate. In conclusion, VTC representations, in contrast to neural networks, fail to veridically represent objects when visual appearance is dissociated from animacy, probably due to a biased processing of visual features typical of animate objects.

## Introduction

A fundamental goal in visual neuroscience is to reach a deep understanding of the neural code underlying object representations; how does the brain represent objects we perceive around us? Over the years, research has characterized object representations in the primate brain in terms of their content for a wide range of visual and semantic object properties such as shape, size, or animacy (Konkle and Oliva, 2012; Nasr et al., 2014; Bracci and Op de Beeck, 2016; Kalfas et al., 2017). More recently, our understanding of these multidimensional object representations has been lifted to a higher level by the advent of so-called deep-convolutional neural networks (DNNs) that do not only reach human behavioral performance in image categorization (Russakovsky et al., 2014; He et al., 2015; Kheradpisheh et al., 2016a), but also appear to develop representations that share many of the properties of primate object representations (Cadieu et al., 2014; Guclu and van Gerven, 2014; Khaligh-Razavi and Kriegeskorte, 2014; Yamins et al., 2014; Guclu and van Gerven, 2015; Kubilius et al., 2016). This correspondence extends to the representation of several object dimensions, such as shape properties (Kubilius et al., 2016), and the distinction between animate and inanimate objects (Khaligh-Razavi and Kriegeskorte, 2014). Hence, the availability of computational models that can mimic human recognition behavior and neural information processing in its full complexity offers exciting possibilities for understanding human object vision (Kriegeskorte, 2015). Here, we provide an important test of this similarity between artificial and biological brain representations by focusing upon a particularly challenging organizing principle of human ventral visual cortex: object animacy.

Perception of animacy has played an essential role through the evolution and survival of our species. This animacy bias is evident in both bottom-up perceptually driven contexts (Gao et al., 2009; Gao et al., 2010; Scholl and Gao, 2013), even after controlling for stimulus low-level visual properties (New et al., 2007), as well as more high-level cognitive phenomena such as pareidolia, where – most often – animate objects (faces or animals) are perceived in meaningless, random noise images (e.g., clouds). At the neural level, our visual system includes specific neural mechanisms with selectivity for animate entities, such as animals and humans, relative to inanimate objects (Kanwisher et al., 1997; Downing et al., 2001). What are the dimensions underlying animacy percepts? One proposal suggests that animacy representations in visual cortex reflect the psychological dimension of perceiving something as being a living entity (Caramazza and Shelton, 1998; Tremoulet and Feldman, 2000; Gao et al., 2009). Alternatively, animacy representations might be “not aware” of object animacy per se but instead reflect stimulus visual aspects such as appearance – whether something looks like a living entity or not. Here we test these two alternatives and compare whether the representation of animacy converges across artificial and biological brains.

We explicitly dissociated object appearance (how an object looks like) from object category (what the object really is) – two dimensions that are typically correlated in natural images, to test whether (1) both biological and artificial brains represent animacy in the same way, and whether (2) animacy percepts reflect object appearance or object category. We show that activity patterns in visual cortex were confused by animal appearance: a cow-shaped mug was more similar to a cow as opposed to a mug. In contrast, DNNs correctly categorized a cow-mug as an inanimate object. As a consequence, rather surprisingly, deep neural networks, which were never explicitly trained on animacy, outperform ventral occipitotemporal cortex representations in categorizing objects as being animate or inanimate.

## Materials and Methods

### Participants

The study included 16 adult volunteers (9 males; mean age, 30 years). Informed consent to take part in the fMRI experiment was signed by all participants. The ethics committee of the KU Leuven approved the study.

For the fMRI experiment, due to excessive head motion, all data from one participant was excluded. In addition, one run was excluded in two participants, and two runs were excluded in two other participants. The head motion exclusion criterion was set to +/− 3 mm (equal to 1 voxel size) and defined before data collection. For behavioral ratings, two participants were excluded due to technical problems during data collection.

### Stimuli

The stimulus set included nine different triads (27 stimuli in total), each containing: (1) one animal (e.g., a cow), (2) one object (e.g., a mug), and (3) one lookalike object, which resembled the animal (e.g., a cow-shaped mug; Figure 1A). Critically, to dissociate object appearance from object identity, each stimulus in the lookalike condition was matched to the inanimate objects in terms of object identity, and to the animals in terms of animal appearance. That is, the lookalike and object conditions shared the same object identity, size, and other object properties such as function and usage (e.g., the mug and the cow-mug). At the same time, the lookalike and animal conditions shared animal appearance, but differed in animacy; the cow-mug it an object whereas the cow depicts a living animal. This stimulus set was used to acquire behavioral, deep neural network, and neuroimaging data.

**Figure 1.**
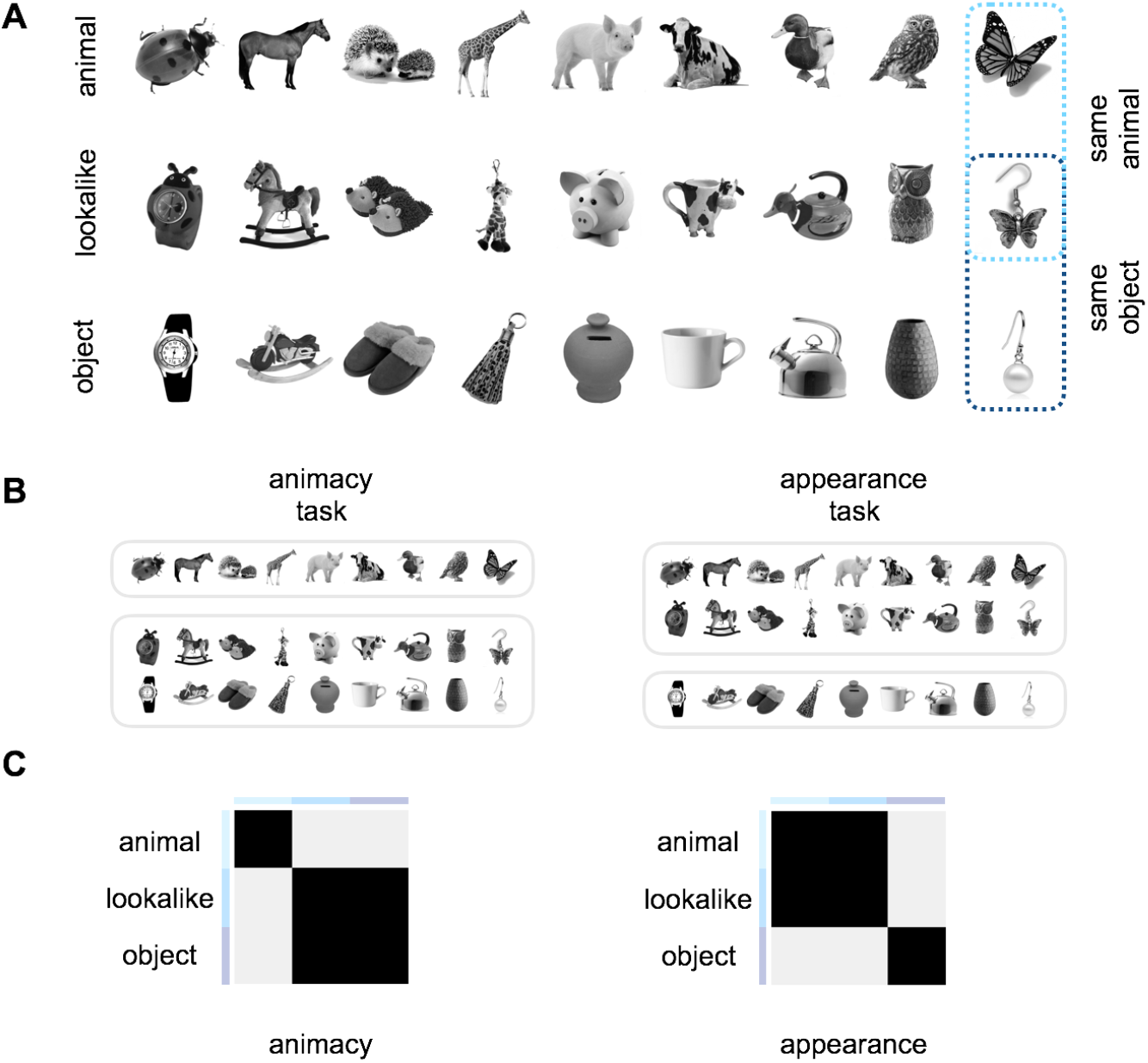
The experimental design. (A) The stimulus set was specifically designed to dissociate object appearance from object identity. We included 9 different object triads. Each triad included an animal (e.g., butterfly), an inanimate object (e.g., earring), and a lookalike object closely matched to the inanimate object in terms of object identity and to the living animal in terms of object appearance (e.g., a butterfly-shaped earring). (B) During fMRI acquisition, participants performed two different tasks counterbalanced across runs. During the *animacy task*, participants judged animacy: “does this image depict a living animal?” During the *animal appearance task*, participants judged animal resemblance: “does this image look like an animal?” Participants responded ‘yes’ or ‘no’ with the index and middle finger. Responses were counterbalanced across runs. (C) Model predictions represent the required response similarity in the two tasks. The animacy model predicts high similarity among images that share semantic living/animate properties, thus predicting all inanimate objects (objects and lookalikes) to cluster together and separately from living animals. On the contrary, the animal appearance model predicts similarities based on visual appearance despite differences in object identity and animacy, thus predicting lookalikes and animals to cluster together and separately from inanimate objects. The two models are independent (r = 0.07).

## Experimental Design

### Behavioral Data

Each participant rated all images (Figure 1A) based on their general similarity. Images were simultaneously rated on a screen within a circle arena following the procedure of Kriegeskorte and Mur (2012). Participants were asked to “arrange the objects according to how similar they are”, thus letting them free to choose what they considered to be the most relevant dimension/s to judge. The object dissimilarity space for behavioral judgments was constructed by averaging results across participants. Only the upper triangle of the dissimilarity matrix was used for subsequent analyses.

### Deep Neural Network Data

To create the object dissimilarity space of our stimulus set, we used the stimuli features that were extracted for the last processing stage (last fully connected layer) of two Deep Neural Networks (DNNs), VGG-19 (Simonyan and Zisserman, 2014) and GoogLeNet (Szegedy et al., 2015). As for behavioral data, only the upper triangle of the dissimilarity matrix was used for subsequent analyses. These DNNs are deep convolutional neural networks, which have been very successful for object recognition tasks in the last few years. They consist of various processing stages, which are often termed as ‘layers’ either individually or in groups, in a feedforward manner that nonlinearly transform an input image volume (width, height, RGB value) into a 1-dimensional vector holding the class scores.

These processing stages include: i) convolutional layers, where a neuron outputs the dot product between a kernel (its weights) and a small region in the input it is connected to, ii) rectified linear unit activation functions, such that the activations of a previous weight layer are thresholded at zero (max(0,x)), iii) max pooling layers performing a downsampling operation along the width and height of the input and lastly, iv) fully connected layers (resembling a multilayer perceptron) that flatten the previous stage’s volume and end up in a I-dimensional vector with the same size as the number of classes. A softmax function is then typically applied to this last fully connected layer’s unit activations to retrieve class probability scores for the classification task. In our experiments, we used the last fully connected layer as the upper layer of the network and not the probability scores from applying the softmax function.

The two networks were trained on ~1.2 million natural images, belonging to 1000 classes for the ImageNet Large Scale Visual Recognition Challenge 2012 (ILSVRC2012). The classes include ~40%animals and ~60% objects, but the networks are not specifically trained to detect animacy. We used the pre-trained models from MatConvNet (http://www.vlfeat.org/matconvnet/)(Vedaldi and Lenc, 2016) toolbox in MATLAB. Before feature extraction, the mean of the ILSVRC2012 training images was subtracted from each image, as to the standards of training both these DNNs. Furthermore, all stimuli were scaled to 224×224 pixels, in accordance with the requirements of the two networks.

VGG-19. This deep neural network from Simonyan and Zisserman (2014) incorporates 16 convolutional layers, in 2 blocks of 2 convolutional layers and 3 blocks of 4 convolutional layers, followed by three fully connected layers. A max pooling operation is applied after each of these five convolutional layer blocks. All, except for the last (fc8), of these weight layers are followed by a RELU function, where the activations are thresholded at zero. A softmax function is then applied to the last fully connected layer (fc8). The top-5 error rate performance of this pre-trained MatConvNet model on the ILSVRC2012 validation data was 9.9%.

GoogLeNet. Google’s entry to ILSVRC2014 (Szegedy et al., 2015). It made use of the ‘Inception’ module, which is a technique used in early versions of Deep Neural Networks for pattern recognition, where a DNN uses several sizes of kernels along with pooling concatenated within one layer, which is similar to integrating all information about parts of the image (size, location, texture etc.). In addition, there is a softmax operation in multiple stages of GoogLeNet, assisting the classification procedure during training along the depth levels of the network. This MatConvNet pretrained model was imported from the Princeton version, not by the Google team, thus there is some difference in performance to other versions probably due to parameter settings during training. The top-5 error rate performance of this model on the ILSVRC2012 validation data was 12.9%.

### fMRI Data

#### Experimental Design

We acquired neuroimaging data by means of an event-related design in two separated sessions, each performed in separated days with no more than 7 days between the first and second session. Each session included six experimental runs as well as additional runs with unrelated stimuli for another experiment (not reported here). The stimuli presentation was controlled by a PC running the Psychophysics Toolbox package (Brainard, 1997) in MATLAB (The MathWorks). Pictures from the stimulus set were projected onto a screen and were viewed through a mirror mounted on the head coil.

Each experimental run (12 in total) lasted 7 min and 14 ms (230 volumes per run). For each subject and for each run, a fully randomized sequence of 27 image trials (repeated 4 times) and 9 fixation trials (repeated 4 times) was presented. Each trial was presented for 1500 ms, followed by a fixation screen for 1500 ms. Each run started and ended with 14 sec of fixation. During the whole experiment, each stimulus was repeated 48 times. While scanning participants performed two different tasks (Figure 1B) counterbalanced across runs. During the animacy task, participants judged animacy (“does this image depict a living animal?”). During the appearance task, participants judged object appearance (“does this image look like an animal?”). Participants responded ‘yes’ or ‘no’ with the index and middle finger. Response-finger associations were counterbalanced across runs.

#### Acquisition Parameters

Imaging data was acquired on a 3T Philips scanner with a 32-channel coil at the Department of Radiology of the University Hospitals Leuven. MRI volumes were collected using echo planar (EPI) T2*-weighted scans. Acquisition parameters were as follows: repetition time (TR) of 2 s, echo time (TE) of 30 ms, flip angle (FA) of 90°, field of view (FoV) of 216 mm, and matrix size of 72 x 72. Each volume comprised 37 axial slices (covering the whole brain) with 3 mm thickness and no gap. The T1 - weighted anatomical images were acquired with an MP-RAGE sequence, with 1×1×1 mm resolution.

#### Preprocessing

Imaging data were preprocessed and analyzed with the Statistical Parametrical Mapping software package (SPM 12, Welcome Department of Cognitive Neurology) and MATLAB. Before statistical analysis, functional images underwent a standard preprocessing procedure to align, co-register and normalize to an MNI (Montreal Neurological Institute) template. Functional images were spatially smoothed by convolution of a Gaussian kernel of 4 mm full width at half-maximum (Op de Beeck, 2010). For each participant, a general linear model (GLM) for both tasks was created to model the 27 conditions of interest and the 6 motion correction parameters (x, y, z for translation and for rotation). Each predictor’s time course was modelled for 3 sec (stimulus presentation + fixation) by a boxcar function convolved with the canonical hemodynamic response function in SPM. We also analyzed data for each task separately, for which purpose a general linear model (GLM) was created for each task separately.

#### Regions of Interest (ROIs) definition

ROIs were defined at the group level with a combination of functional and anatomical criteria. First, we selected all visually active voxels (all stimuli versus baseline) that exceeded the statistical uncorrected threshold p < 0.001. Subsequently, we selected all spatially continuous voxels within anatomical ROIs defined with the Neuromorphometrics atlas in SPM. The following ROIs were defined: V1, and ventro-temporal cortex (VTC) divided into its posterior (post VTC) and anterior (ant VTC) sectors. These ROIs were chosen based on the important role played by VTC in object representation and categorization (Grill-Spector and Weiner, 2014).

### Statistical Analyses

Multivariate analyses were used to investigate the extent to which the two experimental dimensions (object animacy and object appearance) explain representational content in behavioral, DNNs, and brain data (this latter under different task conditions). For statistical tests, we took the following approach. For DNNs models, given that only one similarity matrix for each model was available, statistical significance in the different analyses (e.g., representational similarity analysis) was tested across stimuli with permutation tests (Figure 2) or pairwise t-tests (Figure 3). To be consistent, this approach was also applied to the behavioral data (i.e., these two datasets were directly compared in Figure 2 and 3). For neural data, instead, given that individual subject data was available, we tested significance across subjects with ANOVAs and pairwise t-tests (e.g., Figure 4). Whereas corrections for multiple comparisons were applied for all analyses, for transparency, throughout the text uncorrected p-values are reported. However, when p-values did not survive correction, we noted it in the text. For all statistical tests we report exact p-values up to p = 0.00001, for lower p-values we report p < 0.00001.

**Figure 2.**
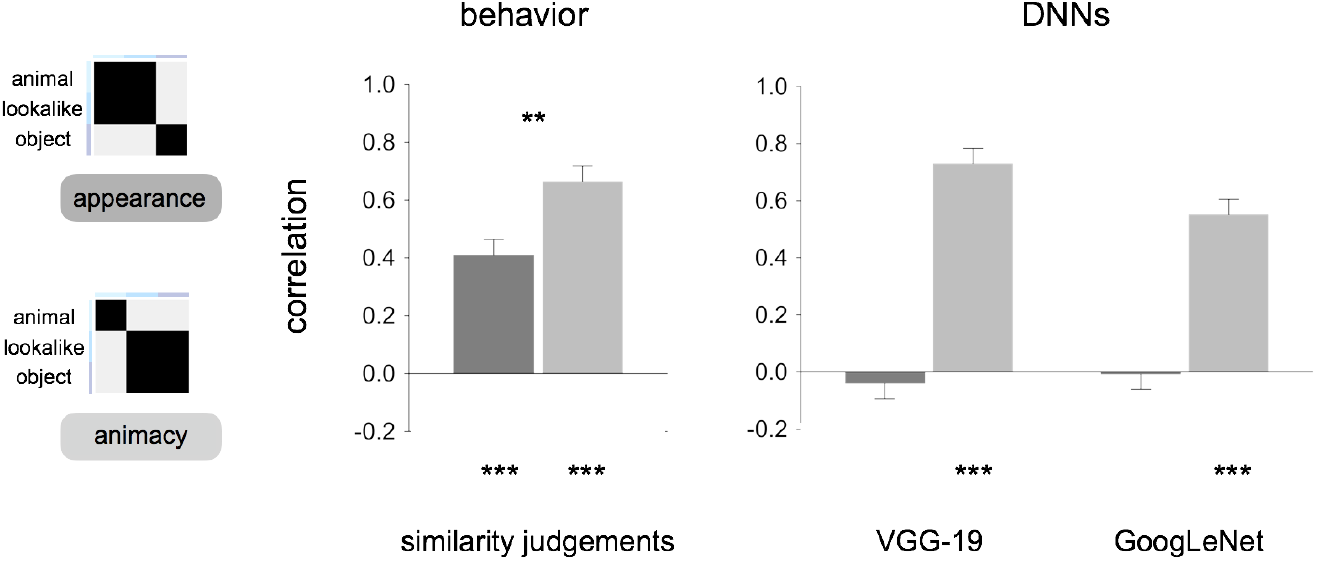
DNNs predict human perception of object animacy but not appearance. Representational similarity analysis (RSA; Kriegeskorte et al., 2008a) results for the appearance model (dark gray) and the animacy model (light gray) are shown for human judgments (left panel), and DNNs (right panel). Asterisks indicate significant values computed with permutation tests (10,000 randomizations of stimulus labels), and error bars indicate SE computed by bootstrap resampling of the stimuli. *** p < 0.0001, ** p < 0.001, * p < 0.01.

**Figure 3.**
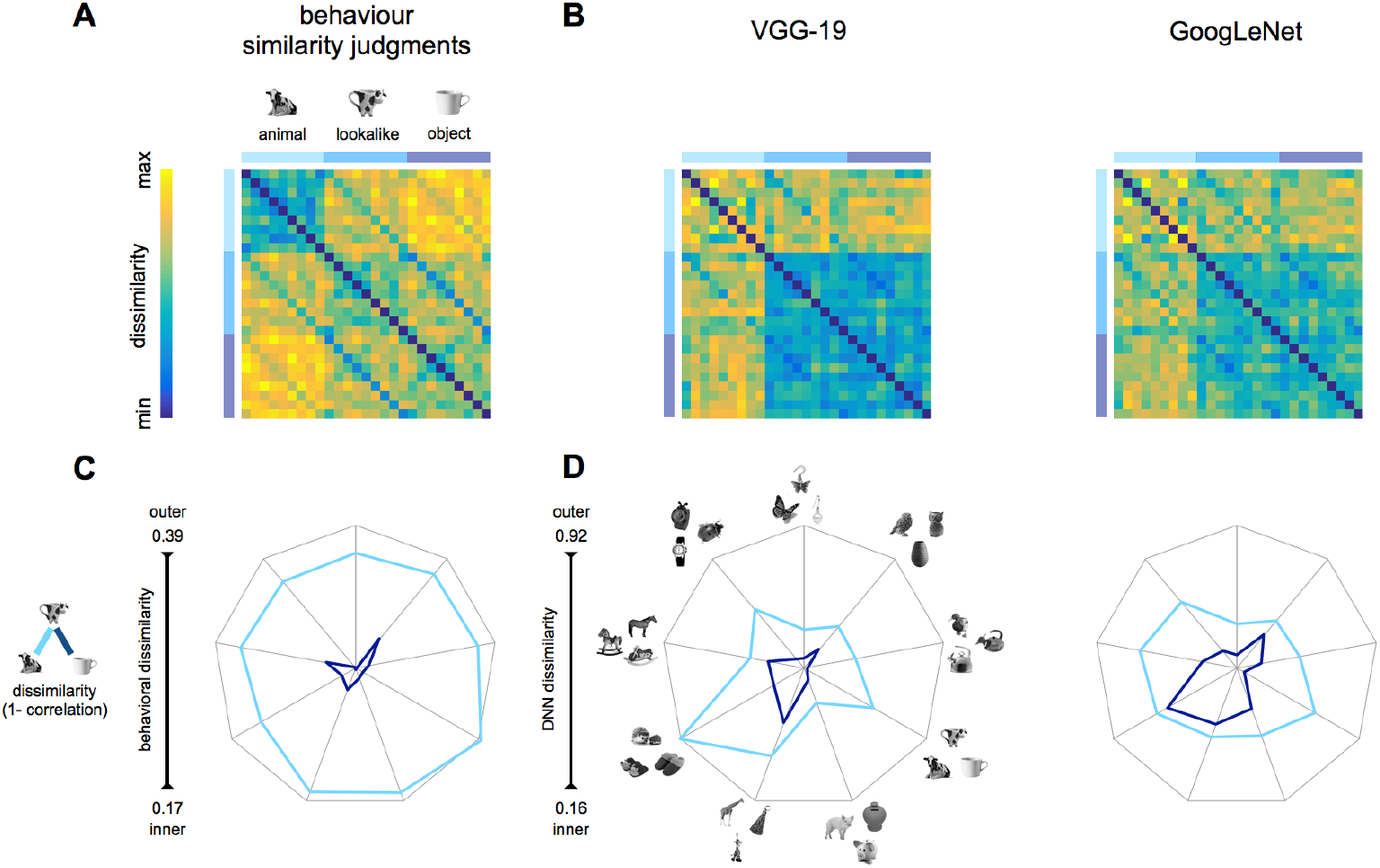
Deep neural networks and human perception predict a representation based on object animacy, over appearance. (A-B) Dissimilarities matrices (upper panel) show stimuli pairwise distances for behavioral judgments (A) and DNNs (B). (C-D) The appearance distance (light blue: animals and lookalikes) and the animacy distance (dark blue: objects and lookalikes) are shown for each object triad (n = 9), for human similarity judgments (C), and DNNs (D). Values are shown on segments, each indicating an object triad. The length of the segments reflects the whole dissimilarity space. Thus, for each chart, the center and the outer points represent the smaller and the largest dissimilarity value, respectively. The closer the value is to the center, the smaller the distance between stimuli within a pair.

**Figure 4.**
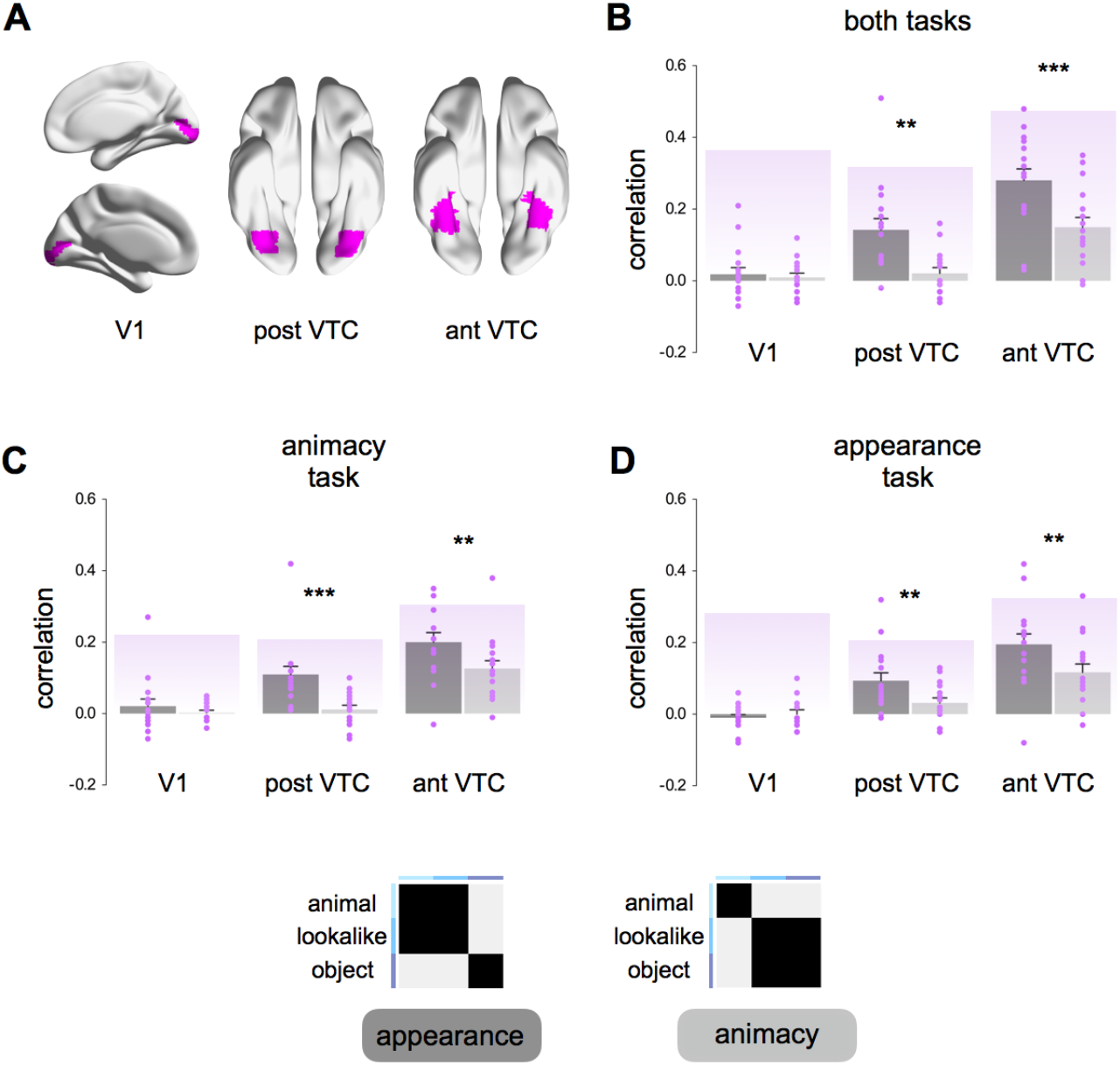
Animal appearance better explains representational content in human visual cortex. A) Group averaged ROIs (V1, post VTC, ant VTC) are shown on an inflated human brain template in BrainNet Viewer (Xia et al., 2013). (B-D) Representational similarity analysis (RSA; Kriegeskorte et al., 2008) results for the appearance model (dark gray) and the animacy model (light gray) are shown for the data combined across the two tasks (B), and for each task separately (C-D). Individual participant’s (n = 16) correlation values are shown in purple. Purple-shaded backgrounds represent reliability values of the correlational patterns taking into account the noise in the data (Materials and Methods). These values give an estimate of the highest correlation we can expect in each ROI. Error bars indicate SEM. Asterisks indicate significant difference between the two models (*** p < 0.001; ** p < 0.01).

#### ROI-based RSA

As before (Op de Beeck et al., 2010), for each voxel within a given ROI, parameter estimates for each condition (relative to baseline) were extracted for each participant and each run and normalized by subtracting the mean response across all conditions. Subsequently, the data set was divided 100 times into 2 random subsets of runs (set-1 and set-2) and the voxel response patterns for each object pair were correlated across these independent data sets. Correlations were averaged across the 100 iterations, thus resulting in an asymmetric 27 × 27 correlation matrix for each task, participant and ROI. For each correlation matrix, cells above and below the diagonal were averaged and only the upper triangle of the resulting symmetric matrix was used in the subsequent analyses (Ritchie et al., 2017). Correlation matrices were converted into dissimilarities matrices (1 minus Pearson’s r) and used as neural input for subsequent analyses. Resulting correlations were Fisher transformed {0.5*log[(1 + r)/(1 − r)]} and tested with ANOVAs and pairwise t-tests.

For each ROI we computed an estimate of the reliability of the dissimilarity matrices (Op de Beeck et al., 2008), which indicates the highest expected correlation in a region given its signal-to-noise ratio. For each subject and each ROI, the 27 × 27 correlation matrix was correlated with the averaged correlation matrix of the remaining participants. The resulting correlation values (averaged across participants) capture noise inherent to a single subject as well as noise caused by inter-subject variability.

#### Whole-brain RSA

In addition to ROI-based RSA, we performed a whole-brain RSA correlating the two models (appearance, animacy) with neural patterns throughout the brain. The whole-brain RSA performed using the volume-based searching approach (Kriegeskorte et al., 2006) was implemented with CoSMo MVPA (Oosterhof et al., 2016). Parameter estimates for each condition (relative to baseline) were extracted for each participant and each run and normalized by subtracting the mean response across all conditions. Resulting values were then averaged across all runs. For each voxel in the brain, a searchlight was defined using a spherical neighborhood with a variable radius, including the 100 voxels nearest to the center voxel. For each searchlight, the neural dissimilarity matrix was computed for the 27 stimuli. The neural dissimilarity matrix (upper triangle) was then correlated with the dissimilarity matrices derived from the two predictive models (Figure 1C). The output correlation values were Fisher transformed and assigned to the center voxel of the sphere. Resulting whole-brain correlation maps for each of the models were directly contrasted and differences were tested using random-effects whole-brain group analysis and corrected with the Threshold-Free Cluster Enhancement (TFCE; Smith and Nichols, 2009) method. Voxel-wise corrected statistical maps (z = 1.64; p < 0.05, one-sided t-test) are highlighted with dotted lines were displayed on a brain template by means of BrainNet Viewer (Xia et al., 2013).

### Data and software availability

All types of brain images and statistics data are available from the authors upon request. The data used for final statistics are made available through the Open Science Framework.

## Results

We constructed a stimulus set to intentionally separate object appearance from object identity, including: 9 animals (e.g., a cow), 9 objects (e.g., a mug), and 9 lookalike objects, which consisted of objects (e.g., a cow-mug) that were matched to the inanimate objects in terms of object identity, and to the animals in terms of appearance. This resulted in a stimulus set of 9 closely matched triads -- 27 stimuli in total (Figure 1A). The mug and the cow-mug represent the same inanimate object, identical in many respects (e.g., function, size, material, and manipulability) but their appearance. Conversely, relative to the cow-mug, the cow is a living animal and despite shared visual features typical of animals, such as eyes, mouth and limbs, differs in all the above semantic and functional properties. With this stimulus design, we dissociate the conceptual animate-inanimate distinction from visual appearance (Fig. 1C).

### Deep neural networks and human perception privilege object animacy over appearance

To test whether DNNs predict human perception on object animacy and appearance, we compared similarity judgments on the stimulus set (Materials and Methods) to the stimulus representation for two recent DNNs (VGG-19: Simonyan and Zisserman, 2014; and GoogLeNet: Szegedy et al., 2015) that were chosen based on their human-like performance in object categorization (Russakovsky et al., 2014; He et al., 2015; Kheradpisheh et al., 2016b). The object dissimilarity for each image pair (1 minus Pearson’s r) was computed for human similarity ratings (Materials and Methods), and for the output vectors of the DNNs’ last fully connected layer, and resulting dissimilarity matrices were correlated with the two independent models. Results revealed similarities but also differences between human judgments and DNNs representations (Figure 2). A significant positive correlation of both models (appearance: r =0.41; p < 0.0001; animacy: r= 0.66; p < 0.0001) with behavioral judgments showed that participants perceived both dimensions, with a significant bias for the animacy percept (p = 0.0006). This bias was particularly strong in DNNs’ representations, with a highly significant correlation with the animacy model (VGG-19: r = 0.73; p < 0.0001; GoogLeNet: r = 0.55; p < 0.0001), but no significant correlation with the appearance model (VGG-19: r = -0.04; GoogLeNet: r = -0.01; p > 0.5, for both DNNs). These results are also evident when inspecting dissimilarity matrices and comparing them to the model matrices (Figure 3 A); animals appear to be separated from objects and lookalikes in both behavioral and DNNs dissimilarity matrices. In addition, animals cluster together in behavioral data with some additional similarities to the lookalike objects, which was absent in DNNs representations.

Despite being orthogonal (r=0.07), both models predict animals and objects being separated from each other. Therefore, in the next analysis we directly test whether lookalikes are represented more similar to animals or other inanimate objects by computing two distance measures (Figure 3C-D): (1) the *appearance* distance (light blue), reflects correlation distance between each animal (e.g., cow) and its matching lookalike object (e.g., cow-mug), and (2) the *animate* distance (dark blue), reflects correlation distance between each object (e.g., mug) and its matching lookalike object (cow-mug).

Confirming the above results, human judgments privilege animacy over object appearance; a test across all object triads showed that the appearance distance (mean = 0.36, SEM = 0.001) was significantly larger than the object distance (mean = 0.019, SEM = 0.001; t_(8)_ = 18.73, p < 0.00001; Figure 3C). Similarly, DNNs are not deceived by animal appearance and represent a cow-mug as being more similar to a mug (animacy distance: VGG-19: mean 0.27, SEM 0.03; GoogLeNet: mean 0.35, SEM 0.05) as oppose to a living cow (appearance distance: VGG-19: mean 0.27, SEM 0.03, t_(8)_ = 4.93, p = 0.001; GoogLeNet: mean 0.35, SEM 0.05, t_(8)_ = 4.65, p = 0.002; Figure 3D). Taken together, these data show that both humans and DNNs set apart animate from inanimate objects in accordance with one of the most reported divisions in ventral occipitotemporal cortex (VTC) (Kriegeskorte et al., 2008b; Grill-Spector and Weiner, 2014).

### Animal appearance explains representational content in the human visual cortex

The above results, from DNNs and human behavior, point to the animacy model as the representational structure to sustain object representations even with a stimulus set that sets object animacy apart from object visual appearance. Based on previous studies, we would expect this organizational principle to have its neural correlate in VTC (Kriegeskorte et al., 2008b; Grill-Spector and Weiner, 2014), possibly accompanied by a further effect of visual appearance (Bracci and Op de Beeck, 2016). We collected human functional neuroimaging (fMRI) data in an event-related design (Materials and Methods). All 27 images were presented in a random order while participants performed two orthogonal tasks (matching the models) counterbalanced across runs (Figure 1B). During the *animacy task*, participants judged whether the image on the screen depicted a living animal (yes or no). During the *appearance task*, participants judged whether the image on the screen looked like an animal (yes or no). In this way, we forced participants to group the lookalike condition in two different ways depending on their properties (Figure 1C): either similarly to the object condition (as in the animacy model) or similarly to the animal condition (as in the appearance model). Thus, in addition to test the two independent predictive models, we can assess any task-related modulation.

We correlated the dissimilarity values predicted by the two models with the dissimilarity matrices derived from neural activity patterns elicited in target regions of interest (ROIs; Figure 4A) in visual cortex. Our main ROI was VTC, divided into its anterior portion (ant VTC) and posterior portion (post VTC). In addition, we included VI as a control ROI. Analyses were performed separately for each task (Figure 4 C-D), but when no task-related effects were observed, we mostly focus on results for the combined dataset.

To investigate the role of animacy and animal appearance in driving the VTC organization, correlation values were tested in a 3×2 ANOVA with ROI (V1, post VTC, ant VTC) and Model (appearance, animacy) as within-subject factors. Results revealed a significant ROI x Model interaction (F_(2,30)_ = 9.40, p = 0.001; Figure 4B), thus highlighting differences in the relation between the two models and representational content in the three ROIs. No positive correlations were found in V1 (lookalike: t <1; animacy: t < 1), suggesting that our stimulus set was constructed appropriately to investigate neural representations without trivial confounds with low-level visual features. Unexpectedly, and differently from DNNs and behavioral results, neural representations in anterior and posterior VTC were significantly more correlated with the appearance model than the animacy model (post VTC: t_(15)_ = 3.85, p = 0.002; ant VTC: t_(15)_ = 4.00, p = 0.001). In addition, we also observed differences between the anterior and posterior portion of VTC. Whereas in anterior VTC both models were significantly correlated with the neural data (appearance: t_(15)_ = 8.70, p < 0.00001; animacy: t_(15)_ = 5.36, p < 0.00001), in posterior VTC, only correlations with the appearance model significantly differed from baseline (lookalike: t_(15)_ = 4.56, p = 0.0004; animacy: t < 1.5). Replicating results within subjects, data analyzed for the two tasks separately did not reveal any task effect in VTC (Figure 4C-D). In both tasks, the appearance model was significantly more correlated with the neural data as oppose to the animacy model in post VTC (animacy task: t_(15)_ = 3.83, p = 0.002; appearance task : t_(15)_ = 3.05, p = 0.008) and ant VTC (animacy task: t_(15)_ = 2.97, p = 0.009; appearance task: t_(15)_ = 3.44, p = 0.004). Furthermore, none of the ROIs showed an effect of task in a direct statistical comparison (ant VTC: F < 1; post VTC: F < 1). To visualize the representational structure in each ROI in more detail, dissimilarities matrices (averaged across tasks) are shown in Figure 6A.

Findings from the ROI-based analysis were backed up by a whole-brain searchlight analysis, which showed a significant preference for the appearance model over the animacy model in (right) VTC (p < 0.05, one-tail t-test, TFCE corrected; Figure 5). The opposite contrast (animacy > appearance) revealed a nonsignificant trend in the occipito-parietal cortex.

**Figure 5.**
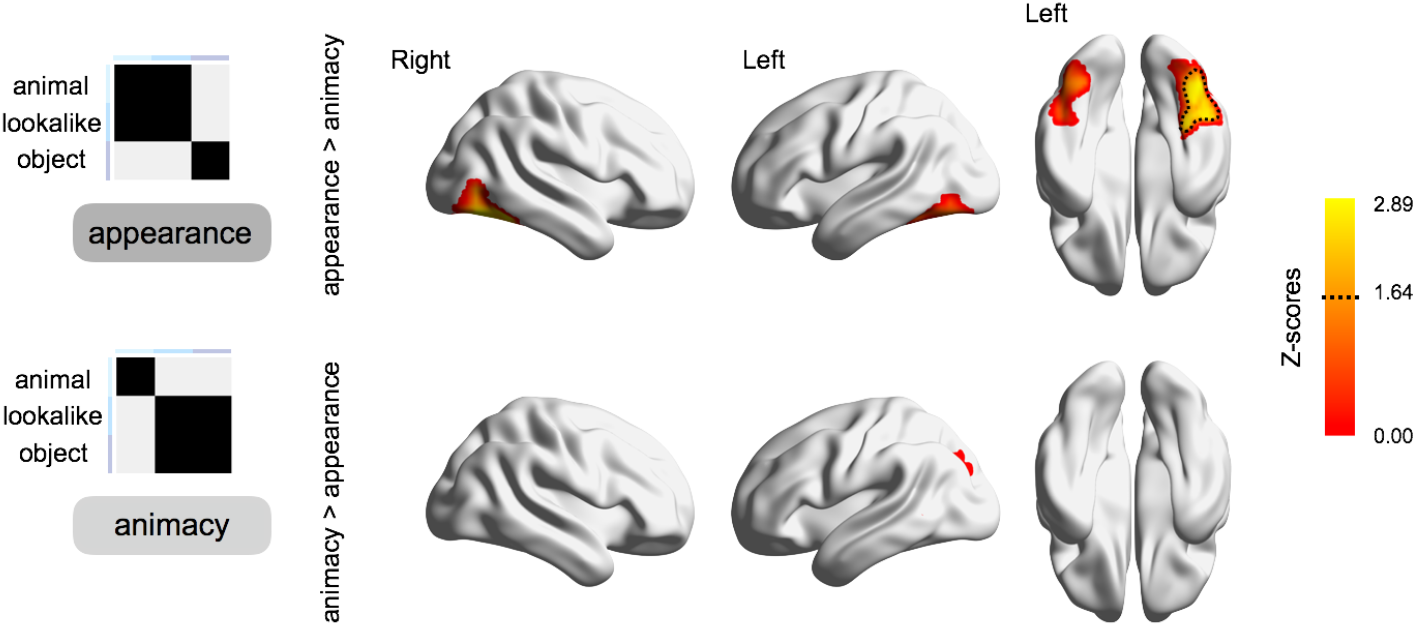
Whole-brain representational similarity analysis. Results of random-effects whole-brain RSA, for the direct contrast between the two predictive models (appearance and animacy) corrected with the Threshold-Free Cluster Enhancement (TFCE; Smith and Nichols, 2009) method. Voxel-wise corrected statistical maps (z = 1.64; p < 0.05, one-sided t-test) are highlighted with dotted lines and displayed on a brain template by means of BrainNet Viewer (Xia et al., 2013).

To further quantify these results, as for behavior and DNNs data, we computed the appearance distance (light blue) and the animacy distance (dark blue) for each object triad. A test across subjects showed that in anterior VTC (Figure 6B), across all triads, the appearance distance (between each animal and its matching lookalike) was significantly smaller (mean: 0.94, SEM: 0.02, t_(15)_ = 3.97, p = 0.001) relative to the animacy distance (between each object and its matching lookalike; mean: 1.02, SEM: 0.01). Converging results where observed when the data were analyzed for the two tasks separately (animacy task: t_(15)_ = 2.62, p = 0.019; appearance task: t_(15)_ = 3.78, p = 0.002), thus highlighting the robustness of these findings. Results did not reach significance in posterior VTC (animacy distance: mean 1.0, SEM: 0.02; appearance distance: mean 0.94, SEM: 0.02; t_(15)_ = 1.94, p = 0.07). Instead, in V1 we observed the opposite trend – smaller distance between neural patterns for lookalikes and objects (animacy distance: mean 0.52, SEM: 0.01; appearance distance: mean 1.01, SEM: 0.01; t_(15)_ = 4.81, p = 0.0002); thus, suggesting that results in ant VTC cannot be explained by confounds between appearance and low-level visual properties.

**Figure 6.**
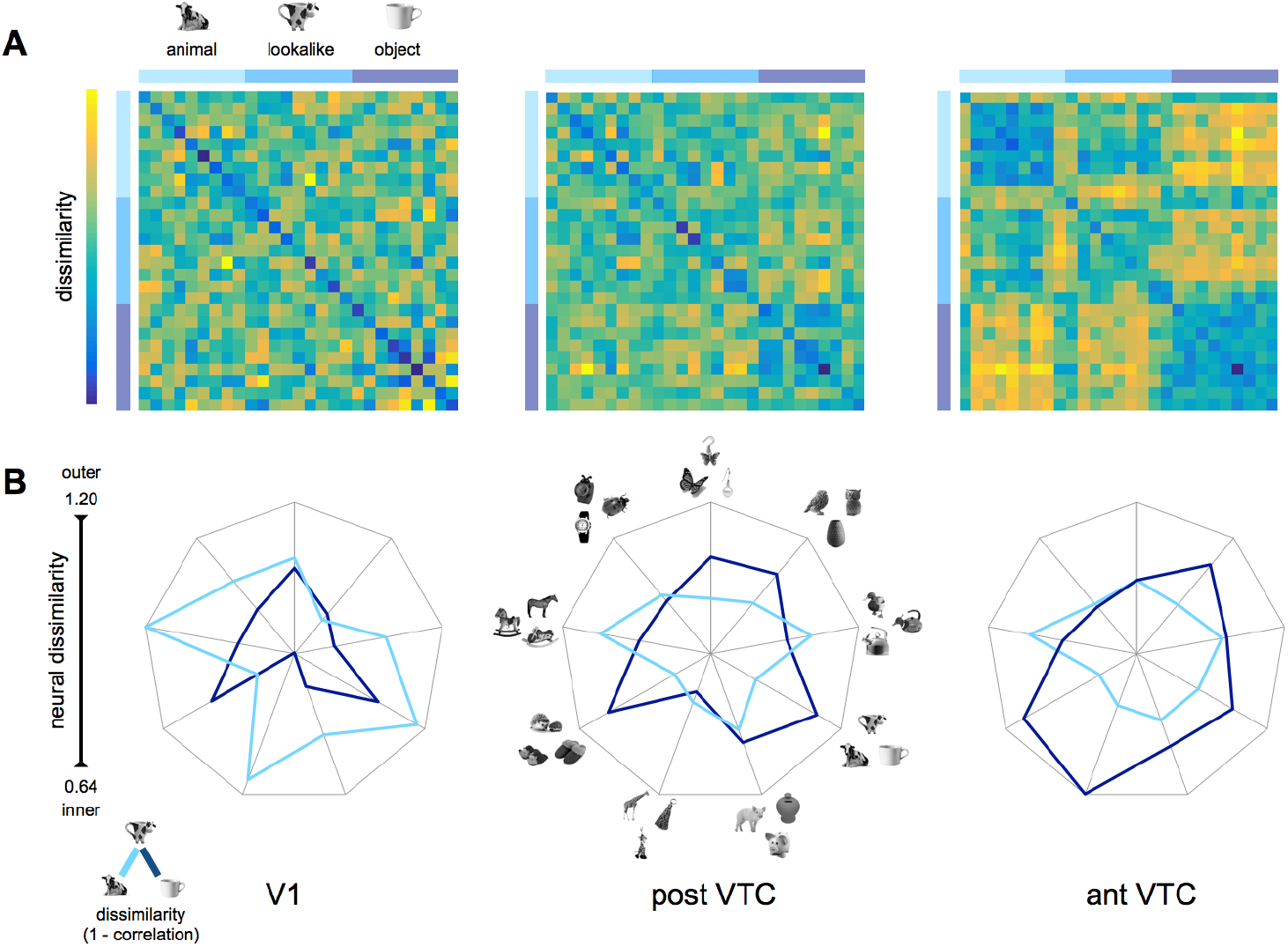
The neural similarity space in VTC reflects animal appearance. (A) The neural dissimilarity matrices (1 minus Person’s r) derived from the neural data (averaged across subjects and tasks) show pairwise dissimilarities among stimuli in the three ROIs (data averaged across tasks). (B) For the three ROIs, the appearance distance (light blue: between animals and lookalikes) and the animacy distance (dark blue: between objects and lookalikes) are shown for each object triad (n = 9). Values are averaged across subjects (n = 16) and tasks (animacy and appearance) and shown on segments, each referring to an object triad. The length of the segments reflects the whole dissimilarity space. Thus, for each chart, the inner and outer points represent the smaller and the largest dissimilarity value, respectively. The closer values are to the center, the smaller the distance between stimuli within pairs.

Together these results show that the animacy organization reported in VTC is largely explained by animal appearance rather than object animacy per se. Inanimate objects (e.g., cow-mug) that share with animals neither functional properties nor animacy, are represented closer to living animals than to other inanimate objects with which they share object category (e.g., a mug), and functional/semantic properties. The animal appearance might relate to high-level visual features such as faces, eyes, limbs, and bodies, which are not generally present in inanimate objects. This result, replicated across tasks, is particularly striking in light of the aforementioned results from human judgments and DNNs, which privilege object animacy over appearance.

### Do VTC representations distinguish between lookalikes and real animals?

In anterior VTC the two (independent) models both differ from baseline. Therefore, although the lookalikes were closer to animals than to objects, lookalikes and animals might be encoded separately in VTC. Another possibility is that the positive correlation with the animacy model is fully driven by the distinction between animals and (non-lookalike) objects (which both predictive models share), without any representation of the lookalikes as being different from real animals (on which the two predictive models make opposite predictions). In other words, does VTC discriminate between animals and objects that look like animals or is it largely blind to this category distinction? If VTC does not distinguish between animals and objects that have animal appearance then there should be no difference between within- and between-condition correlations for these two categories. In what follows we investigated this hypothesis.

To this aim, we computed the *category index*, which reflects the extent to which representations for two conditions can be discriminated from each other. That is, for each subject and condition, the average within-condition correlation (e.g., comparing an animal with other animals) and between-condition correlation (e.g., comparing animals with lookalikes or objects) were calculated. For each condition pair (i.e., animal-lookalike, animal-object, and lookalike-object), the *category index* was computed as follows: first averaging the two within-condition correlations (for animals and lookalikes) and then subtracting the between-condition correlation for the same two categories (animals-lookalikes). For between-condition correlations, diagonal values (e.g., duck and duck-kettle) were excluded from this computation to avoid intrinsic bias between conditions that share either animal appearance (i.e., duck and duck-kettle) or object category (i.e., kettle and duck-kettle). Category indexes above baseline indicate that two category representations are separated. In both VTC regions, the category index could distinguish between animals and objects (ant VTC: t_(15)_ = 6.00, p = 0.00002; post VTC: t_(15)_ = 4.03, p = 0.001; Figure 7, left panel), and objects and lookalikes (ant VTC: t_(15)_ = 7.40, p < 0.00001; post VTC: t_(15)_ = 4.02, p = 0.001). The category index between animals and lookalikes was significant in ant VTC (t_(15)_ = 5.38, p = 0.00007) but did not differ from baseline (after correcting for multiple comparisons: p 0.05/9 = p < 0.005) in post VTC (t_(15)_ = 2.34, p = 0.03). Furthermore, in both ROIs, the category index for animals and lookalikes was significantly smaller than the other two category indexes (ant VTC: t_(15)_ > 3.49, p < 0.004, for both indexes; post VTC: t_(15)_ > 3.45, p < 0.005, for both indexes). In V1 none of the conditions could be distinguished (t < 1, for all condition pairs). As for previous analyses, these results were replicated when data were analyzed for each task separately (the category index for animals vs. lookalikes was significantly smaller than the other two indexes: animacy task: p < 0.004; appearance task: p < 0.01). Thus, VTC representations reflect animal appearance much more than animacy, with a small remaining difference between animals and lookalikes in the anterior part of VTC (but not posterior VTC).

**Figure 7.**
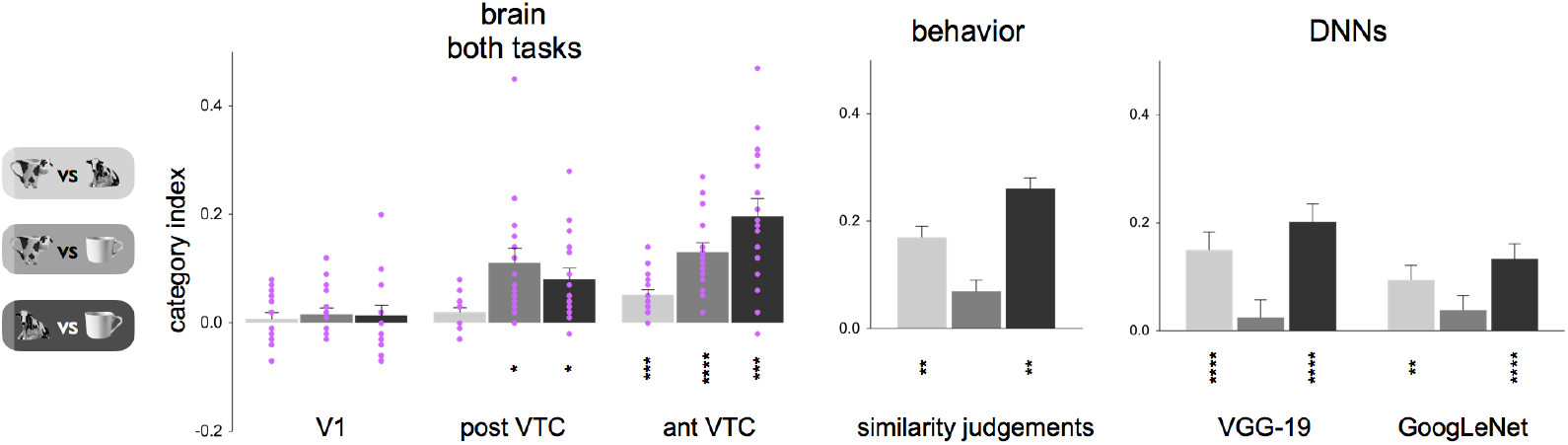
VTC representations differ in category discriminability from DNNs and human behavior. The *category index* reflects representational discriminability among the three stimulus categories (animals, lookalikes, and objects) and is computed for each condition pair by subtracting the average of between-condition correlations (e.g., for animals and lookalikes), from the average of within-condition correlations (e.g., for animals and lookalikes). Results are reported for neural data (left panel), behavior (middle panel), and DNNs (right panel). Light gray: animals vs. lookalikes; gray: lookalikes vs. objects; dark gray: animals vs. objects. For neural data, individual participant’s (n = 16) data points are shown in purple. Asterisks indicate significant values relative to baseline, and error bars indicate SEM. For behavioral and DNNs data, asterisks indicate significant values relative to baseline, computed with permutation tests (10,000 randomizations of stimulus labels), and error bars indicate SE computed by bootstrap resampling of the stimuli. **** p < 0.00001, *** < p < 0.0001, ** < p < 0.001, * < p < 0.01.

Different results were observed for behavioral judgments and DNNs. In both cases, category indexes distinguished between animals and objects (behavior: p = 0.0001; VGG-19: p < 0.00001; GoogLeNet: p < 0.00001), between animals and lookalikes (behavior: p = 0.0006; VGG-19: p < 0.00001; GoogLeNet: p = 0.0005), but did not differ between lookalikes and objects (behavior: p = 0.03 – num. comparisons: p 0.05/3 = p < 0.01; VGG-19: p = 0.46; GoogLeNet: p = 0.15). This suggests that differently from VTC representations, human judgments and convolutional neural network take object category into account; two mugs belong to the same object category regardless of their shape.

Together, these results show that representations in VTC are not predicted either by convolutional neural networks or human judgments. Whereas the former privileges the role of animal appearance, the latter favours the role of superordinate object category such as objects versus animals.

### The coding for animal appearance in VTC interferes with invariant object identification

Previous studies have shown that VTC contains information about object identity, with a high degree of invariance for a variety of image transformations (Gross et al., 1972; Desimone et al., 1984; Grill-Spector et al., 1998; Grill-Spector et al., 1999). Despite the reported finding that a lookalike object is represented very differently from other objects, VTC representations might still allow for object identity identification (e.g. recognizing that a cow-mug is a mug). To allow this, the representation of a lookalike object should be more similar to another object from the same basic category (e.g., a cow-mug and a regular mug) than to other objects. We operationalize this as the prediction that the within-triad correlation between each lookalike and its corresponding non-lookalike object would be significantly higher than the average of correlation of this same lookalike to objects from other triads. We call this the *identity index*. That is, for each subject and for each lookalike object, we took the on-diagonal correlation (e.g., between the cow-mug and the mug) and subtracted the average of off-diagonal correlations (e.g., between the cow-mug and the remaining objects). We computed the *identity index* separately for animals *(animal identity index:* is a cow-mug closer to a cow relative to other animals?) and objects *(object identity index:* is a cow-mug closer to a mug relative to other objects?). Figure 8 shows results for the dataset combined across the two tasks.

**Figure 8.**
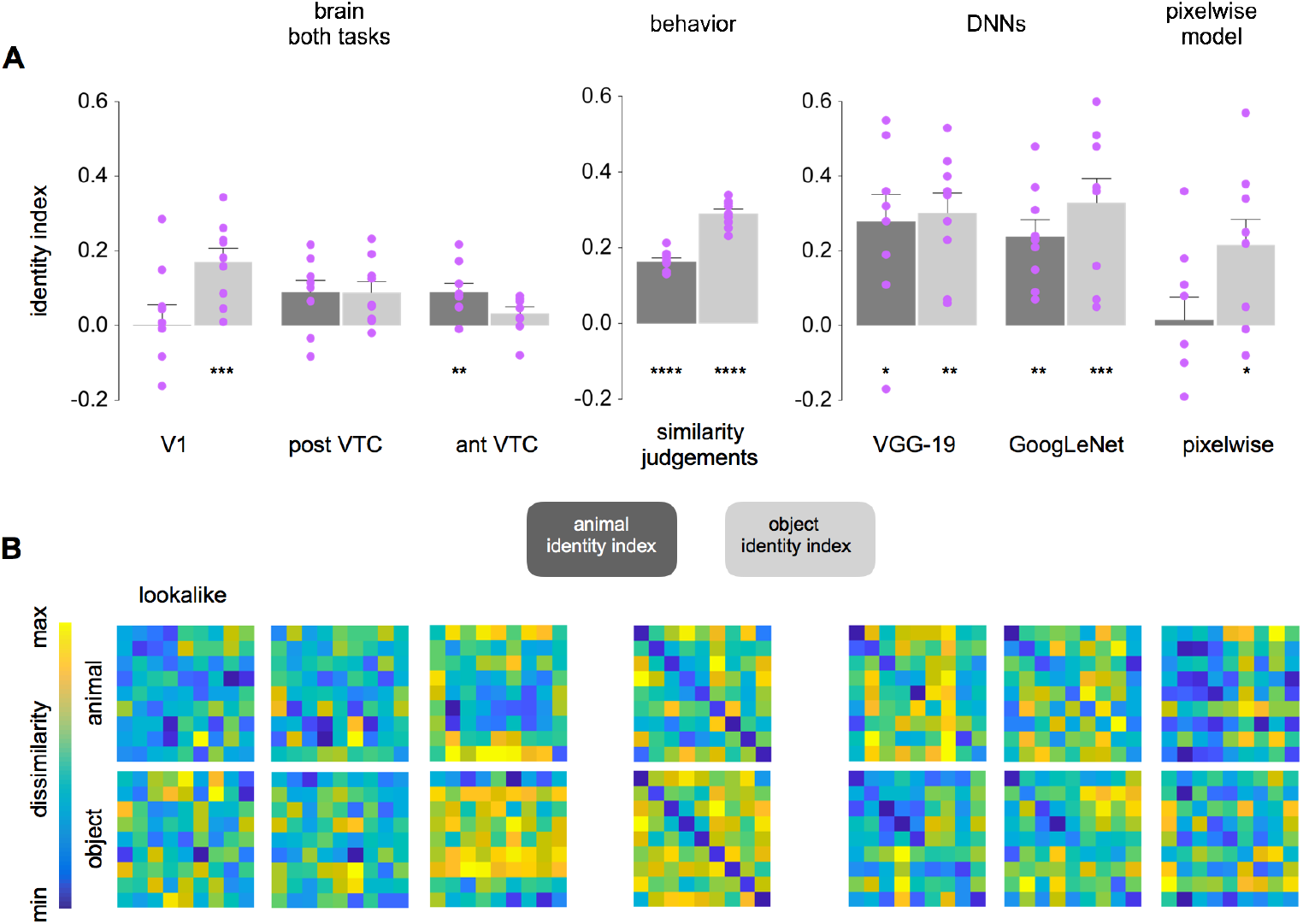
VTC representations sustain animal identity, but not object identity, categorization **(A)** The *identity index* reflects information for individual object ad animal pairs (e.g., the cow-mug and the mug represent the same object; the cow-mug and the cow represent the same animal) and is computed separately for each condition (animals and objects). For each lookalike object (n= 9) we took the on-diagonal correlation (e.g., between the cow-mug and the mug) and subtracted the average off-diagonal correlations (e.g., between the cow-mug and the remaining objects). The identity index for animals and objects was computed for the brain data (V1, post VTC, and ant VTC), behavioral data (similarity judgments), DNNs (VGG-19, GoogLeNet), and the image pixelwise data. Light gray: animal identity index; dark gray: object identity index. Asterisks (**** p < 0.00001, *** < p < 0.0001, ** < p < 0.001, * p < 0.01) indicate significant values relative to baseline, and error bars indicate SEM. (B) For each dataset, dissimilarity matrices used to compute the identity index are shown separately for animals and objects.

For brain data, a test across all conditions (Figure 8, left panel) revealed differences in the amount of stimulus identity information carried in the three ROIs. In V1 there was significant identity information for objects (t_(8)_ = 4.76, p = 0.001) but not for animals (t < 1). This result can be explained considering differences in image low-level visual properties across conditions; objects and lookalikes were more similar to each other relative to animals and lookalikes. Confirming this interpretation, the same trend was observed for image pixel-wise information (Figure 8A, right panel), where we observed significant identity information for objects (t_(8)_ = 3.16, p = 0.01) but not for animals (t < 1). In posterior VTC, neither identity index survived correction for multiple comparisons (p 0.05/6 = p < 0.008; animals: t_(8)_ = 2.82, p = 0.02; objects t_(8)_ = 3.09, p = 0.01). In anterior VTC object identity information decreased and animal identity information increased; here only the animal identity index was significantly above baseline (animals: t_(8)_ = 3.88, p = 0.005; objects: t_(8)_ = 1.93, p = 0.09). Results were replicated when data were analyzed separately for the two tasks (ant VTC: animals, p < 0.009; objects, p > 0.4, for both tasks). Together, these results suggest that representational content in anterior VTC is differently biased to represent animal and object’s identity, containing more information for the former.

In line with previous analyses, different results were observed for behavioral judgments and DNNs. For human similarity judgments, a test across conditions revealed significant identity information for both animals (t_(8)_ = 17.98, p < 0.00001) and objects (t_(8)_ = 24.44, p < 0.00001; Figure 8A, middle panel). Similarly, DNNs were able to discriminate individual stimulus identities for animals (VGG-19: t_(8)_= 3.84, p = 0.005; GoogLeNet: t_(8)_ = 5.42, p = 0.0006) as well as objects (VGG-19: t_(8)_ = 5.67, p = 0.0005; GoogLeNet: t_(8)_ = 5.04, p = 0.001; Figure 8A, right panel).

Strikingly, after years of searching for models that would match the tolerance in the most advanced category representations in primate ventral visual cortex, we have now found an object transformation for which neural networks sustain invariant object recognition to a higher degree than anterior VTC representations.

## Discussion

With a stimulus set that specifically dissociates object appearance from object category, we investigated the characteristics of the previously reported organization of object representations in the superordinate distinction of animate versus inanimate in the human brain, deep neural networks, and behavior. Human behavioral judgments and neural networks privilege animacy over appearance. Conversely, representations in ventral occipitotemporal cortex mostly reflect object appearance. Thus, although deep neural networks can largely predict human behavior, representations in ventral occipitotemporal cortex deviate from behavior and neural network representations.

Our results can be summarized as follows. First, representational content in VTC reflects animal appearance more than object identity and animacy; even though the mug and the cow-mug share many high-level properties (e.g., object identity, size, function, manipulation), as well as low-level properties (e.g., shape and texture), VTC represents a cow-mug closer to a real cow than to a mug. Second, VTC representations are not explained by either human perception or DNNs, which were not deceived by object appearance, and judged a cow-mug being closer to a mug as opposed to a real cow. Third, given its bias in favor of animal appearance, VTC representations are remarkably poor in providing information about object identity, that is, to reflect that a cow-mug is a mug. This is not a desirable property for a ‘what’ pathway that, according to uniformly held views in visual neuroscience (e.g., DiCarlo et al., 2012), builds up representations that sustain reliable and transformation-invariant object identification and categorization. Fourth, VTC representations are not modulated by task demand, and the similarity in responses to animals and lookalike objects persists even when participants performed a task requiring focusing on object animacy.

The animacy division is considered one of the main organizational principles in visual cortex (e.g., Grill-Spector and Weiner, 2014), but information content underlying this division is highly debated (Baldassi et al., 2013; Grill-Spector and Weiner, 2014; Nasr et al., 2014; Bracci and Op de Beeck, 2016; Bracci et al., 2017b; Kalfas et al., 2017). Animacy and other category distinctions are often correlated with a range of low- and higher-level visual features such as the spatial frequency spectrum (Nasr et al., 2014; Rice et al., 2014) and shape (Cohen et al., 2014; Jozwik et al., 2016), but the animacy structure remains even when dissociated from such features (Bracci and Op de Beeck, 2016). Here we question what we imply with animacy. Previous reports suggest that the extent to which an object is perceived as being alive and animate is reflected in VTC representations, giving rise to a continuum where those animals perceived more animate (e.g., primates) are closely represented to humans, whereas those animals perceived less animate (e.g., bugs) are represented away from humans and closer to inanimate objects (Connolly et al., 2012; Sha et al., 2015). Contrary to this prediction, our results suggest that information content underlying the animacy organization does not relate to the animacy concept: whether an object is perceived as being alive and animate is close to irrelevant. Instead, what mostly matters is animal appearance; that is, whether an inanimate object lookalikes and shares high-level visual features with animals. Indeed, in VTC, inanimate objects such as a mug, a water kettle, or a pair of slippers with animal features (e.g., eyes, mouth, tail) are represented close to living animals (Figure 4,6).

Does this mean that it is all about animal appearance? Probably not: our results showed that VTC representational content can be explained by both models, though uncorrelated (r = 0.07; Figure 4), and carries enough information to distinguish between real animals and lookalikes (Figure 7). What we suggest is that information about animals in VTC is overrepresented relative to information about objects – to the extent that even inanimate objects, if having animal appearance, are represented similarly to animate entities. A VTC bias towards animal representations was further supported by results showing significant information for animal’s identity, which was not observed for objects; that is, VTC representations contain information to discriminate that a duck-shaped water kettle represents a duck, but not to discriminate that it is a kettle (Figure 8). An everyday life example of this bias is our strong predisposition to perceive faces (and animals) in meaningless patterns such as toast or clouds (pareidolia), which results in face-like activations in the fusiform face area (Hadjikhani et al., 2009). To our knowledge, similar effects reported for inanimate objects are rare. We could speculate that having a system devoted to detecting animate entities and biased to represent similarly animals and inanimate objects that resemble animals, might have been beneficial from an evolutionary perspective. The chance for our ancestors to survive in the wild necessitated fast and successful detection of predators. Such a biased system would have been advantageous: better mistaking a bush for a lion than the other way around. Thus, the VTC bias to represent lookalikes similarly to real animals might be the result of an evolution-long “training”, similarly to the way, in artificial intelligence, neural networks, trained to recognize animals, start to detect animals in random-noise images (Szegedy et al., 2015).

This speculation points to one possible explanation for a particularly surprising aspect of our findings, namely the discrepancy in representations between VTC at one hand and artificial DNN at the other hand. Indeed, whereas DNNs were able to predict aspects of human perception such as setting apart animals from inanimate objects (Figure 6), and discriminating both animal and object identity (Figure 8), contrary to recent findings (Cadieu et al., 2014; Guclu and van Gerven, 2014; Khaligh-Razavi and Kriegeskorte, 2014; Yamins et al., 2014) our results show that DNNs representations (at the higher processing layers) do not predict human visual cortex representations. Here it is noteworthy that the training history of the DNNs was focused upon the categorization of a large number of categories and was not specifically biased towards e.g. animate objects. This training history might explain why DNNs are very good, and even better than human visual cortex, at identifying a cow-mug as being a mug – not a cow.

Our results show that, across all analyses, human behavior was better predicted by DNNs than human VTC, privileging what an object is (two mugs), as opposed to its visual appearance. Information from DNNs and behavioral judgments did not show a category distinction between lookalikes and inanimate objects (e.g., all objects; Figure 7), yet allowed object identification across changes in appearance (e.g., a cow-mug is a mug; Figure 8). These results differ from previous studies reporting some degree of correspondence between behavior and visual cortex information content (Williams et al., 2007; Carlson et al., 2014; van Bergen et al., 2015). This discrepancy might relate to the choice of task used to obtain the behavioral results that were compared to neural data; here we used a cognitive task (similarity judgments), while a categorization task based on reaction times (e.g., visual search task) might have been a better predictor for visual cortex representations (Proklova et al., 2016; Cohen et al., 2017).

Finally, visual cortex representations were not modulated by tasks. Despite the different focus on object appearance in one case (appearance task), and on object identity in the other case (animacy task), the neural data was striking similar across task sessions revealing no difference in any of the performed analyses. These results confirm, and add to, recent findings showing that task demand appears not to have much influence on the representational structure of ventral visual cortex (Harel et al., 2014; Bracci et al., 2017a; Bugatus et al., 2017; Hebart et al., 2018).

The mere observation that a stimulus that looks like an animal or part of it, such as a face, can elicit animal- or face-like responses in ventral visual cortex is not novel, with studies on face pareidolia being the most relevant example (Hadjikhani et al., 2009). What makes our study novel and our results striking, is the availability of behavioral and neural network data that makes a very different prediction; the systematic comparison and dissociation of neural, behavioral, and DNN representations; the quantification of the degree of bias towards representing appearance rather than animacy; the persistence of this bias in VTC even when performing a task requiring to report animacy; and the detrimental effect of this bias upon the informational content about object identity.

To summarize, these results highlight substantial differences in the way categorical representations in the human brain, human judgments, and deep neural networks characterize objects. Whereas DNN representations resemble human behavior and heavily weight object identity, representations in ventral visual cortex are very much constrained by object appearance. Based on these findings, we might have to reconsider the traditional view of the human visual cortex as a general object recognition machine like the artificial neural networks are trained to be.

## Author Contributions

S.B. and H.O.B designed research; S.B. performed research; S.B. and I.K. analyzed data; S.B., I.K. and H.O.B. wrote the paper.

## Acknowledgements

S.B. was funded by FWO (Fonds Wetenschappelijk Onderzoek) through a postdoctoral fellowship (12S1317N) and a Research Grant (1505518N). H.O.B. was supported by the European Research Council (ERC- 2011-StG-284101), a federal research action (IUAP-P7/11), the KU Leuven Research Council (C14/16/031), and a Hercules grant ZW11_10.

